# Low-Dose Ad26.COV2.S Protection Against SARS-CoV-2 Challenge in Rhesus Macaques

**DOI:** 10.1101/2021.01.27.428380

**Authors:** Xuan He, Abishek Chandrashekar, Roland Zahn, Frank Wegmann, Jingyou Yu, Noe B. Mercado, Katherine McMahan, Amanda J. Martinot, Cesar Piedra-Mora, Sidney Beecy, Sarah Ducat, Ronnie Chamanza, Sietske Rosendahl Huber, Leslie van der Fits, Erica N. Borducchi, Michelle Lifton, Jinyan Liu, Felix Nampanya, Shivani Patel, Lauren Peter, Lisa H. Tostanoski, Laurent Pessaint, Alex Van Ry, Brad Finneyfrock, Jason Velasco, Elyse Teow, Renita Brown, Anthony Cook, Hanne Andersen, Mark G. Lewis, Hanneke Schuitemaker, Dan H. Barouch

## Abstract

We previously reported that a single immunization with an adenovirus serotype 26 (Ad26) vector-based vaccine expressing an optimized SARS-CoV-2 spike (Ad26.COV2.S) protected rhesus macaques against SARS-CoV-2 challenge. In this study, we evaluated the immunogenicity and protective efficacy of reduced doses of Ad26.COV2.S. 30 rhesus macaques were immunized once with 1×10^11^, 5×10^10^, 1.125×10^10^, or 2×10^9^ vp Ad26.COV2.S or sham and were challenged with SARS-CoV-2 by the intranasal and intratracheal routes. Vaccine doses as low as 2×10^9^ vp provided robust protection in bronchoalveolar lavage, whereas doses of 1.125×10^10^ vp were required for protection in nasal swabs. Activated memory B cells as well as binding and neutralizing antibody titers following vaccination correlated with protective efficacy. At suboptimal vaccine doses, viral breakthrough was observed but did not show evidence of virologic, immunologic, histopathologic, or clinical enhancement of disease compared with sham controls. These data demonstrate that a single immunization with a relatively low dose of Ad26.COV2.S effectively protected against SARS-CoV-2 challenge in rhesus macaques. Moreover, our findings show that a higher vaccine dose may be required for protection in the upper respiratory tract compared with the lower respiratory tract.

## Introduction

Immune correlates of protection against SARS-CoV-2 have yet to be defined in humans. We recently reported that purified IgG from convalescent rhesus macaques protected naïve animals against SARS-CoV-2 challenge in a dose-dependent fashion and that cellular immune responses may also contribute to protection^1^. We previously demonstrated that an adenovirus serotype 26 (Ad26) vector^2^ expressing a stabilized SARS-CoV-2 Spike^3,4^, termed Ad26.COV2.S, effectively protected rhesus macaques against SARS-CoV-2 infection and protected hamsters against severe SARS-CoV-2 disease^5,6^. In these studies, vaccine-elicited binding and neutralizing antibodies correlated with protection^5,6^. DNA vaccines, mRNA vaccines, ChAdOx1 vectors, and inactivated virus vaccines have also been reported to protect against SARS-CoV-2 challenge in macaques^7-11^.

In multiple SARS-CoV-2 vaccine studies in nonhuman primates, protection in the upper respiratory tract appeared less robust than protection in the lower respiratory tract^6-8,11^. These data have raised the possibility that protection against asymptomatic infection may be more difficult to achieve than protection against severe pneumonia in humans. However, the role of vaccine dose in protection in the upper and lower respiratory tracts has not previously been defined. Moreover, suboptimal vaccine doses can be utilized to assess the theoretical concern of vaccine-associated enhanced respiratory disease (VAERD), although VAERD has not been reported to date in SARS-CoV-2 vaccine studies in animals or humans.

In this study, we assessed the immunogenicity and protective efficacy of a titration of Ad26.COV2.S dose levels to evaluate immune correlates of protection, to define the role of reduced vaccine doses in protecting different anatomic respiratory compartments, and to assess the possibility of VAERD. We observed that low doses of Ad26.COV2.S protected against Ad26.COV2.S challenge in the lower respiratory tract but that higher vaccine dose levels were required to protect in the upper respiratory tract. Suboptimal vaccine dose levels resulted in reduced protective efficacy, but no evidence of VAERD was observed.

## Results

### Vaccine Immunogenicity with Reduced Ad26.COV2.S Dose Levels

We immunized 30 adult male and female rhesus macaques with a single dose of 1×10^11^, 5×10^10^, 1.125×10^10^, or 2×10^9^ viral particles (vp) Ad26.COV2.S (N=5/group) or sham (N=10) at week 0 (**Fig. S1**). We observed induction of RBD-specific binding antibodies by ELISA in animals that received the 1×10^11^, 5×10^10^, and 1.125×10^10^ vp doses by week 2 and in animals that received the 2×10^9^ vp dose by week 4 (**Fig. 1A**). Neutralizing antibody (NAb) responses were assessed using a pseudovirus neutralization assay^11-13^ and were observed in the majority of animals in the three higher dose groups by week 2, with increasing titers through week 6 (**Fig. 1B**). NAb titers remained low in the 2×10^9^ vp group at weeks 2 and 4 but became detectable in all animals by week 6, suggesting slower kinetics and lower magnitude NAb responses (**Fig. 1B**).

**Figure 1.**
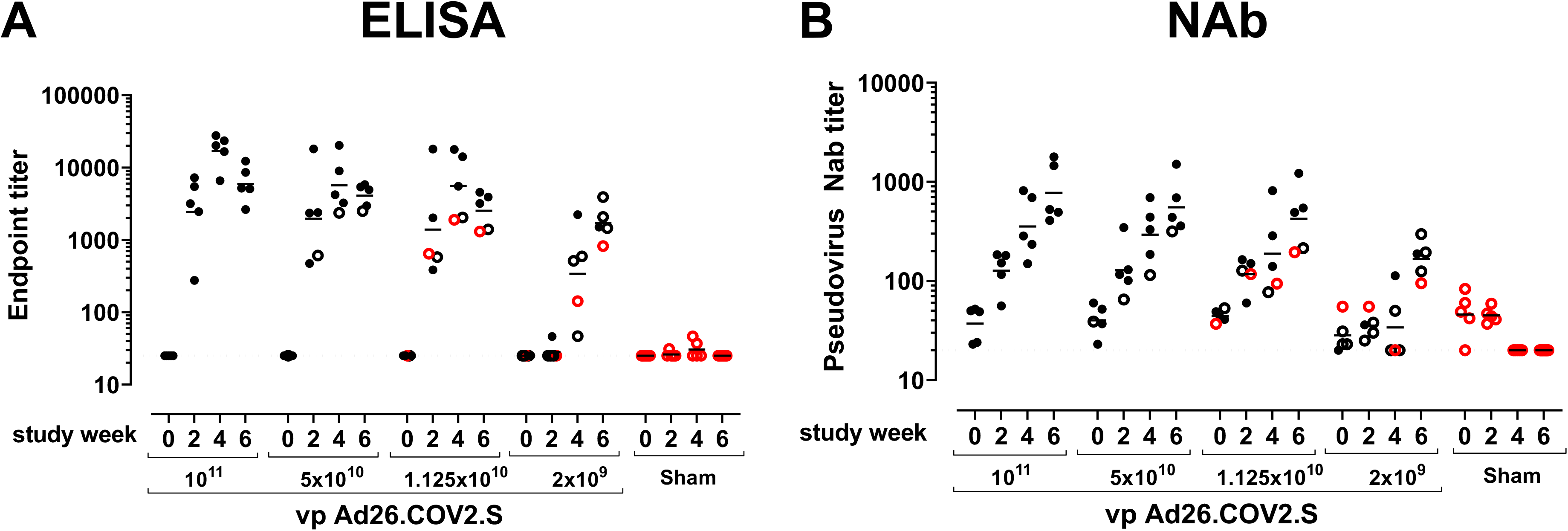
Humoral immune responses in vaccinated rhesus macaques. Humoral immune responses were assessed at weeks 0, 2, 4, and 6 by (**A**) RBD-specific binding antibody ELISA and (**B**) pseudovirus neutralizing antibody (NAb) assays elicited by a single immunization of 1×10^11^, 5×10^10^, 1.125×10^10^, or 2×10^9^ vp Ad26.COV2.S (N=5/group) or sham controls (N=10). Horizontal bars reflect geometric mean responses. Dotted lines reflect assay limit of quantitation. Solid black circles indicate animals that showed no virus in BAL and NS following challenge, open black circles indicate animals that showed virus in NS but not BAL following challenge, and open red circles indicate animals that show virus in both BAL and NS following challenge.

IFN-γ ELISPOT assays using pooled S peptides demonstrated T cell responses in the majority of vaccinated animals that received the 1×10^11^, 5×10^10^, and 1.125×10^10^ vp doses at week 4, although there was a trend for lower responses with lower vaccine doses (**Fig. 2A**). In animals that received the 2×10^9^ vp dose, only 2 of 5 animals developed detectable ELISPOT responses (**Fig. 2A**). IL-4 ELISPOT responses were undetectable (**Fig. 2B**), suggesting induction of Th1-biased responses and consistent with prior findings^6^.

**Figure 2.**
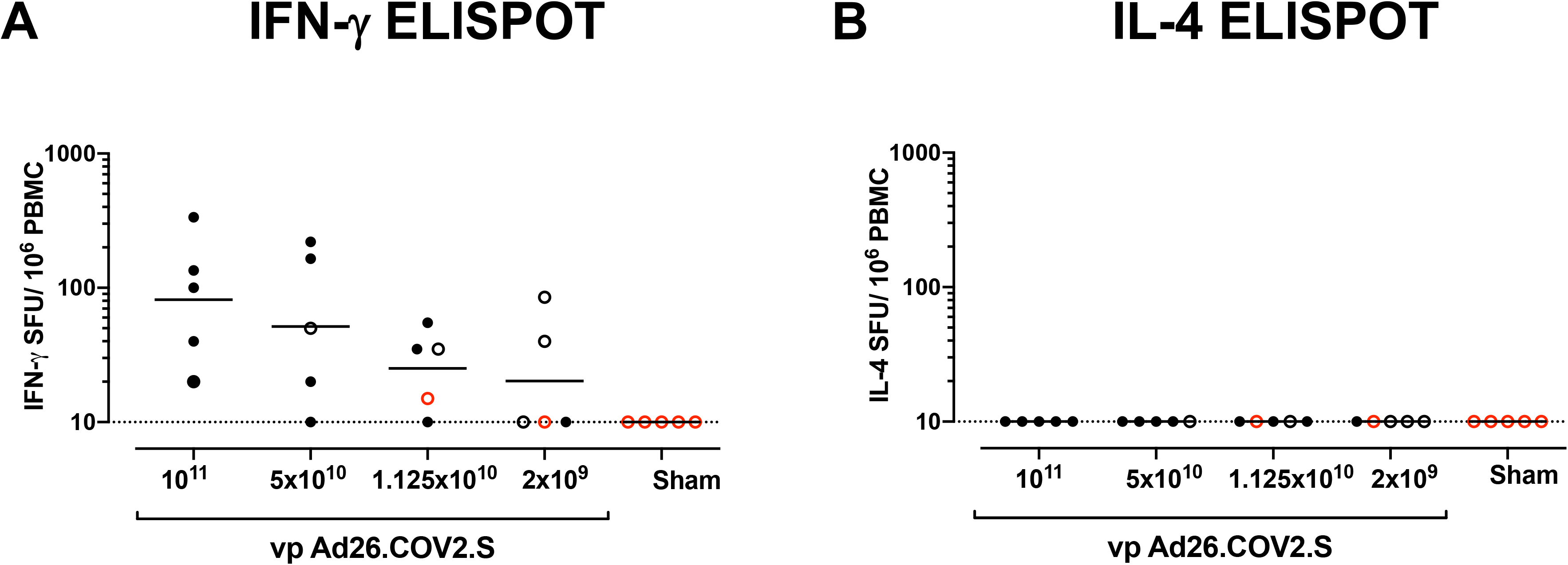
T cell responses in vaccinated rhesus macaques. Cellular immune responses were assessed at week 4 following immunization by (**A**) IFN-γ and (**B**) IL-4 ELISPOT assays in response to pooled S peptides. Horizontal bars reflect geometric mean responses. Dotted lines reflect assay limit of quantitation. Solid black circles indicate animals that showed no virus in BAL and NS following challenge, open black circles indicate animals that showed virus in NS but not BAL following challenge, and open red circles indicate animals that show virus in both BAL and NS following challenge.

We next monitored B cell responses following vaccination by multiparameter flow cytometry. SARS-CoV-2 RBD-specific IgG+ B cells were detected in peripheral blood by week 2 following vaccination and generally expressed the activation marker CD95 and the memory marker CD27 (**Fig. S2**), suggesting activated memory B cells^14-16^. RBD-specific activated memory B cells expanded following vaccination in a dose-dependent fashion, with robust responses in all animals that received 1×10^11^ vp but marginal responses in animals that received 2×10^9^ vp (**Fig. 3**). The frequency of RBD-specific activated memory B cells strongly correlated with NAb and ELISA titers (P<0.0001, R=0.7997 and P<0.0001, R=0.8851, respectively, twosided Spearman rank-correlation tests) and moderately correlated with IFN-γ ELISPOT responses (P=0.0063, R=0.5310) (**Fig. S3**).

**Figure 3.**
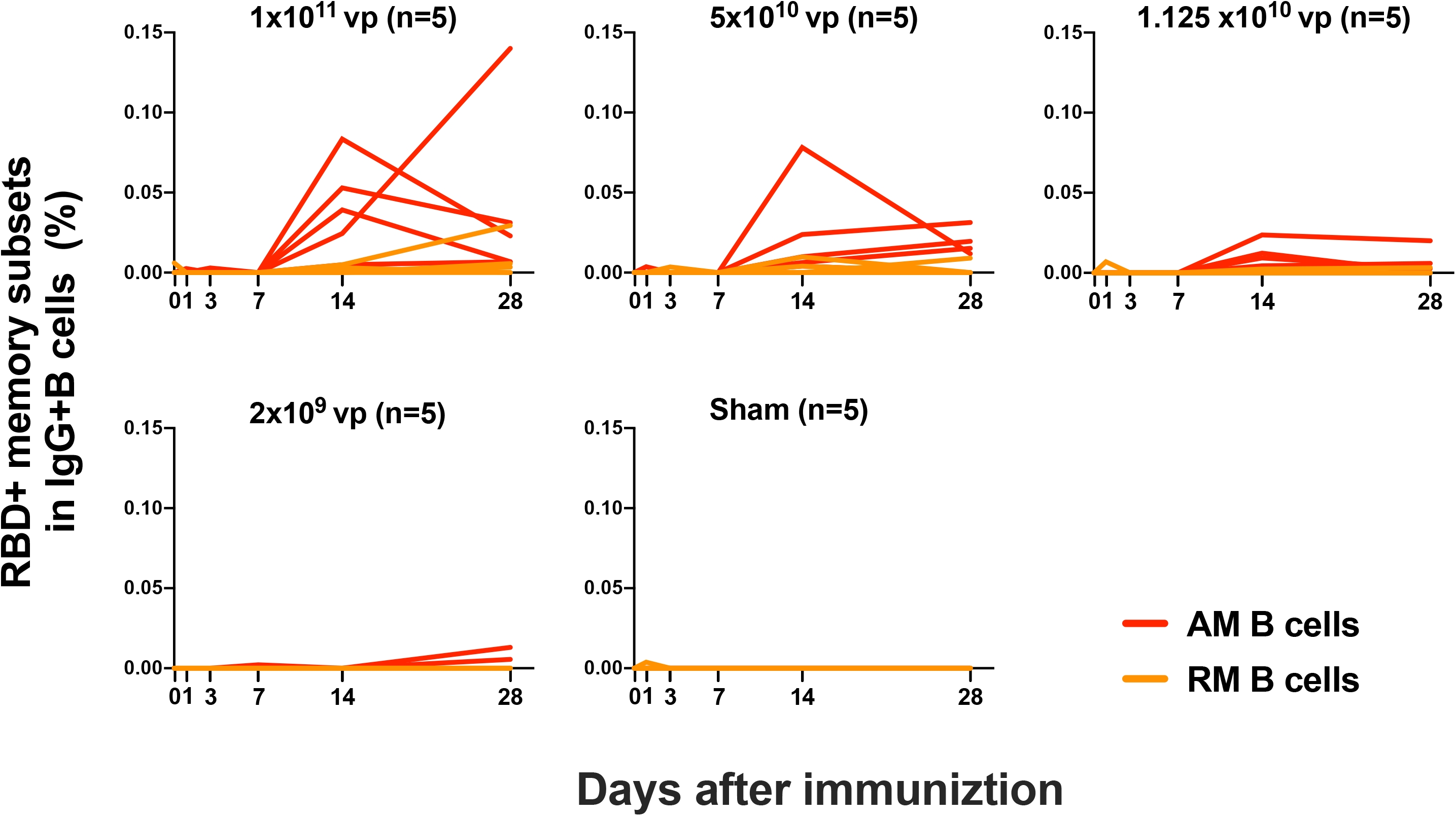
B cell responses in vaccinated rhesus macaques. Frequencies of RBD-specific CD27+CD95+ activated memory B cells (red) and resting memory B cells (orange) in total IgG+ B cell populations on days 0, 1, 3, 7, 14, 28 following Ad26.COV2.S immunization.

### Protective Efficacy with Reduced Ad26.COV2.S Dose Levels

We challenged all animals at week 6 with 1.0×10^5^ TCID50 SARS-CoV-2 by the intranasal (IN) and intratracheal (IT) routes^1,6,11,13^. We assessed viral loads in bronchoalveolar lavage (BAL) and nasal swabs (NS) by RT-PCR specific for subgenomic mRNA (sgRNA), which is believed to measure replicating virus^13,17^. All 10 sham controls were infected and showed a mean peak of 4.45 (range 3.2-6.5) log_10_ sgRNA copies/ml in BAL (**Fig. 4A**). In contrast, vaccinated animals demonstrated no detectable virus in BAL (limit of quantitation 1.69 log_10_ sgRNA copies/ml), with the exception of one animal in the 2×10^9^ vp group and one animal in the 1.125×10^10^ vp group (**Fig. 4A**). Similarly, all sham controls showed a mean peak of 5.68 (range 3.8-6.9) log_10_ sgRNA in NS (**Fig. 4B**). In contrast with limited viral breakthroughs in the BAL, 80% (4 of 5) of animals in the 2×10^9^ vp group, 40% (2 of 5) of animals in the 1.125×10^10^ vp group, and 20% (1 of 5) of animals in the 5×10^10^ vp group showed viral breakthroughs in NS (**Fig. 4B**). These data suggest that a higher vaccine dose may be required for protection in the upper respiratory tract compared with protection in the lower respiratory tract. Suboptimal vaccine dose levels led to loss of protection in NS but did not result in enhanced virus replication compared with the sham controls.

**Figure 4.**
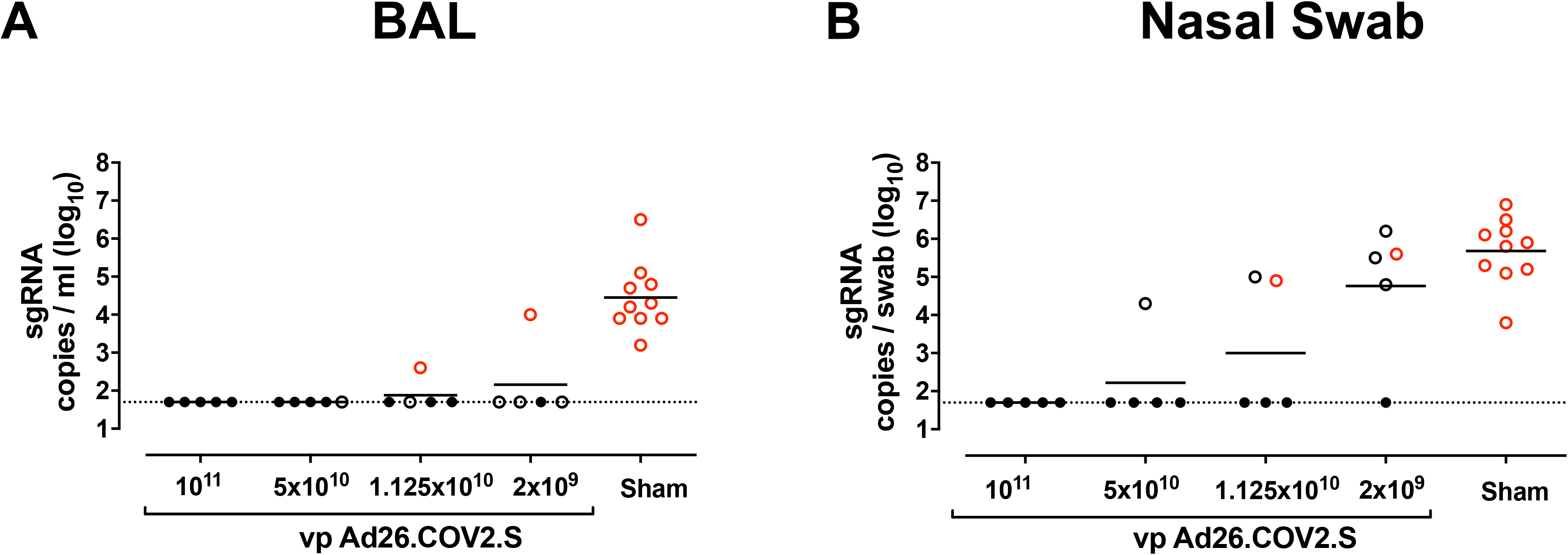
Protective efficacy following SARS-CoV-2 challenge. Rhesus macaques were challenged by the intranasal and intratracheal routes with 1.0×10^5^ TCID50 SARS-CoV-2. (**A**) Peak log_10_ sgRNA copies/ml (limit of quantification 50 copies/ml) were assessed in bronchoalveolar lavage (BAL) following challenge. (**B**) Peak log_10_ sgRNA copies/swab (limit of quantification 50 copies/swab) were assessed in nasal swabs (NS) following challenge. Horizontal lines reflect geometric mean values. Solid black circles indicate animals that showed no virus in BAL and NS following challenge, open black circles indicate animals that showed virus in NS but not BAL following challenge, and open red circles indicate animals that show virus in both BAL and NS following challenge.

The log_10_ ELISA and NAb titers at week 6 inversely correlated with peak log_10_ sgRNA in BAL (P<0.0001, R=-0.8489 and P<0.0001, R=-0.8343, respectively, two-sided Spearman rank-correlation tests; **Fig. 5**) and in NS (P<0.0001, R=-0.7765 and P<0.0001, R=-0.8436, respectively, two-sided Spearman rank-correlation tests; **Fig. 5**). Moreover, the frequency of RBD- and S-specific activated memory B cells inversely correlated with peak log_10_ sgRNA in NS (P<0.0001, R=-0.7196 and P=0.0003, R=-0.6686, respectively, for day 14 responses; P=0.0001, R=-0.6936 and P=0.0014, R=-0.6039, respectively, for day 28 responses; two-sided Spearman rank-correlation tests; **Fig. 6A**). In addition, RBD- and S-specific activated memory B cells were higher in completely protected animals compared with partially protected or nonprotected animals (P=0.0006 and P=0.0005, two-sided Mann–Whitney tests; **Fig. 6B**). Taken together, these data show that both memory B cell responses and binding and neutralizing antibody titers correlated with protection against SARS-CoV-2 in rhesus macaques.

**Figure 5.**
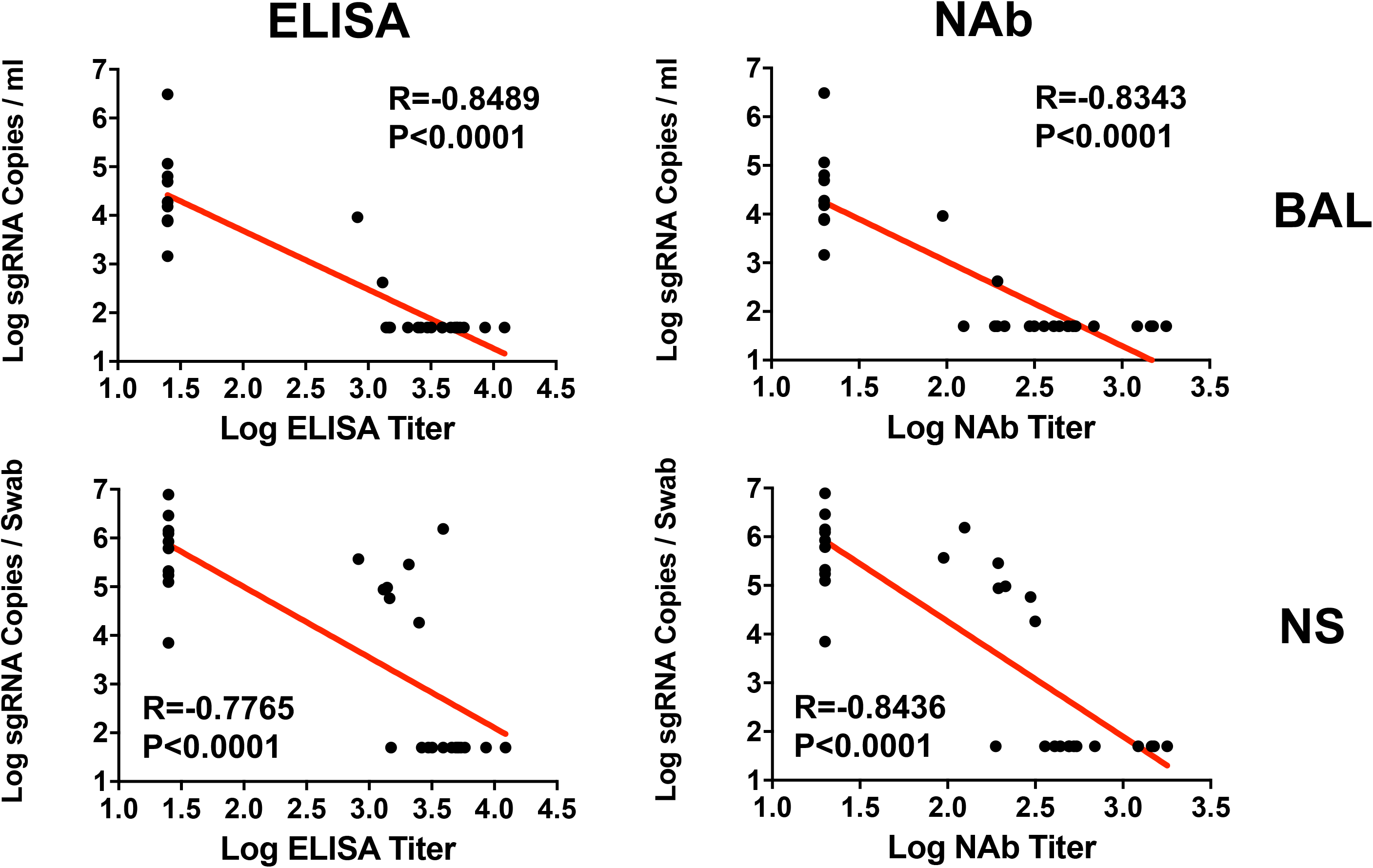
Antibody correlates of protection in BAL and NS. Correlations of log ELISA titers and log NAb titers at week 6 following vaccination with log peak sgRNA copies/ml in BAL and NS following challenge. Red lines reflect the best linear fit relationship between these variables. P and R values reflect two-sided Spearman rank-correlation tests. n=30 biologically independent animals.

**Figure 6.**
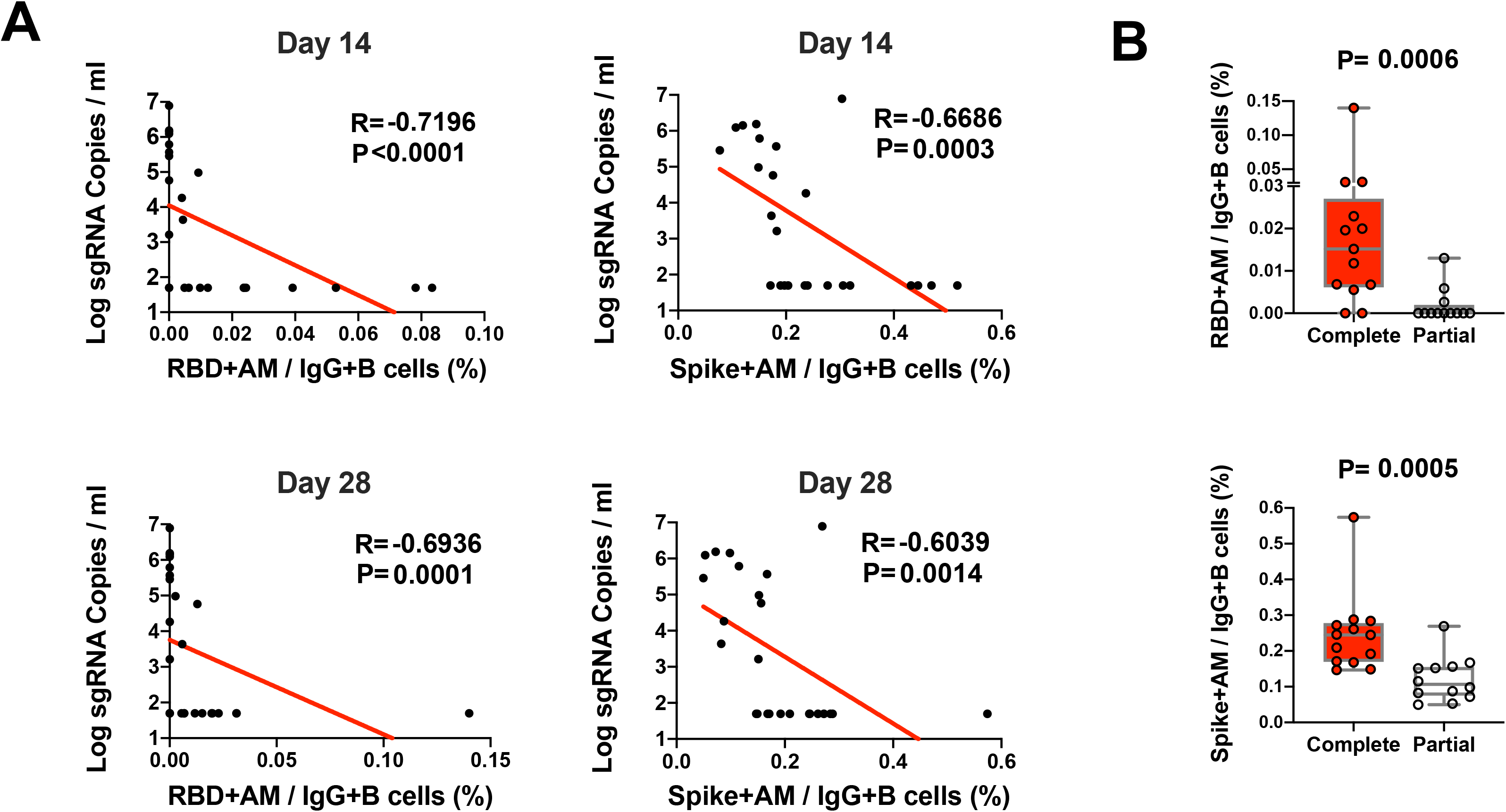
B cell correlates of protection in NS. **(A)** Correlations of RBD- and S-specific activated memory B cell responses at day 14 and day 28 following vaccination with log peak sgRNA copies/ml in NS following challenge. Red lines reflect the best linear fit relationship between these variables. P and R values reflect two-sided Spearman rank-correlation tests. n=25 biologically independent animals. (B) Frequencies of RBD- and S-specific activated memory B cells in completely protected macaques (n=13) and partially protected and non-protected macaques (n=12). P values reflect two-sided Mann–Whitney tests.

### Histopathology Following SARS-CoV-2 Challenge

On day 10 following challenge, animals were necropsied, and lung tissues were assessed by histopathology and immunohistochemistry. We observed focal to locally extensive SARS-CoV-2 associated pathological lesions in sham controls (**Fig. 7, Table 1**). We previously reported histopathologic evidence of viral pneumonia in rhesus macaques on day 2 and day 4 following SARS-CoV-2 infection^13^. On day 10, lungs in sham controls still showed evidence of multifocal interstitial pneumonia with bronchoepithelial syncytia, perivascular mononuclear infiltrates, type II pneumocyte hyperplasia, rare thrombosis, and focal edema and consolidation (**Fig. S4, Table 1**). RNAscope demonstrated in situ hybridization for viral RNA, immunohistochemistry showed staining for SARS nucleocapsid, and infiltrates included Iba-1+ macrophages, CD3+ T cells, and CD20+ B cells (**Fig. S5**). In contrast, vaccinated animals showed minimal histopathologic changes, consistent with background lung pathology, although several animals that received the 2×10^9^ vp dose showed evidence of mild inflammation (**Figs. 7, 8; Table 1**). No evidence of VAERD was observed in animals that received high or suboptimal doses of Ad26.COV2.S, including animals that showed breakthrough viral replication in BAL and/or NS.

**Figure 7.**
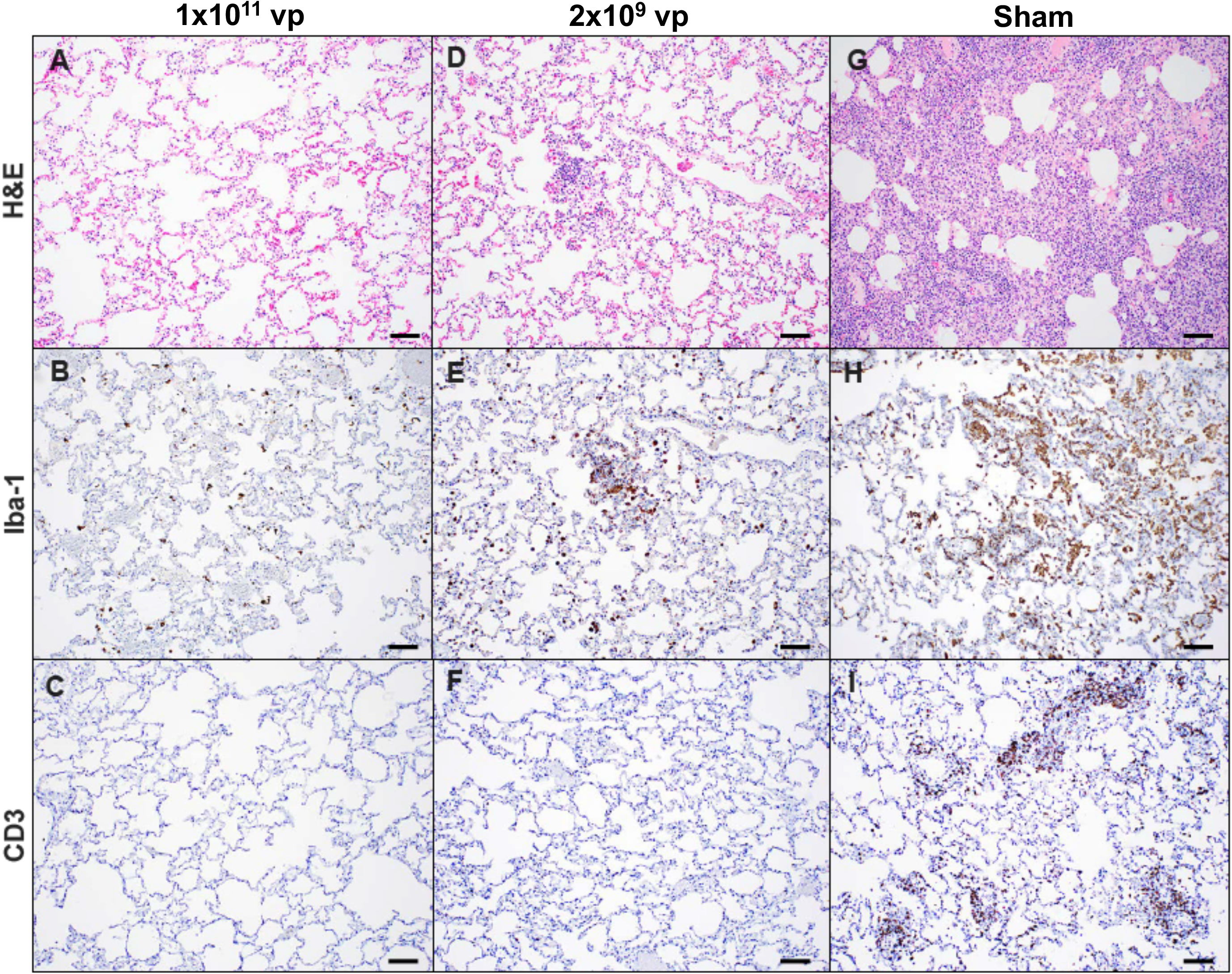
Comparative pathology in vaccinated and unvaccinated animals following SARS-CoV-2 challenge. Representative pathology from animals vaccinated with (**A-C**) 1×10^11^ vp Ad26.COV2.S, (**D-F**) 2×10^9^ vp Ad26.COV2.S, or (**H-I**) sham on day 10 following SARS-CoV-2 challenge. (**A, D, G**) representative H&E histopathology. (**B, E, H**) immunohistochemistry for Iba-1 (macrophages). (**C, G, I**) showing immunohistochemistry for CD3 (T-lymphocytes). Animals that received a high vaccine dose had minimal evidence of SARS CoV-2 pathology (**AC**). Animals that received the lowest vaccine dose showed focal pathology (**D-F**) characterized by increased alveolar macrophages, focal interstitial septal thickening and aggregates of macrophages. Sham vaccinated animals had locally extensive moderate interstitial pneumonia (**G**) characterized by extensive macrophage infiltrates (**H**) and expansion of perivascular and interstitial CD3 T lymphocytes (**I**). Scale bars = 100 microns.

**Table 1:** Histopathology scoring of lung lesions in vaccinated and sham animals following SARS-CoV-2 challenge. Eight lung tissue sections representing one section from each of 7 lung lobes, and two sections from the left cranial lobe, were processed histologically and evaluated microscopically for the histopathological findings listed in the table. The listed histopathological findings were graded on a scale of 0-5, with grade 0 representing no findings, grade 1 minimal histological change, 2: mild, 3: moderate, 4: marked and 5: severe/massive histological change. Histopathology grades were tallied for all lung sections, and average scores for each group were calculated. Total histopathology scores were calculated by adding up scores for the individual pathology parameters. BALT: bronchus-associated lymphoid tissue

| Group | 1 | 2 | 3 | 4 | 5 |
| --- | --- | --- | --- | --- | --- |
| Vaccine | Ad26.COV2.S |  |  |  | Sham |
| Dose (vp) | 1x10<sup>11</sup> | 5x10<sup>10</sup> | 1.125x10<sup>10</sup> | 2x10<sup>9</sup> | NA |
| N | 5 | 5 | 5 | 5 | 10 |
| Median total histopathology score (range) | 10 (9-14) | 11 (9-14) | 11 (8-12) | 12 (9-23) | 30 (16-45) |
| Average total histopathology score (range) | 11 (9-14) | 11 (9-14) | 10 (8-12) | 14 (9-23) | 29 (16-45) |
|  | Lung pathology scores for the individual pathology parameters, average score per group |  |  |  |  |
| Inflammation, interstitial/septal thickening | 1.2 | 2.4 | 2.6 | 2.6 | 7.3 |
| Infiltrates, mononuclear, perivascular | 2.2 | 3.2 | 3.2 | 5.6 | 8.9 |
| Infiltrates, macrophage, alveolar | 4.2 | 4.8 | 3.8 | 4.0 | 6.5 |
| Infiltrates, macrophage, bronchial | 0.2 | 0.4 | 0 | 0 | 1.4 |
| Hyperplasia, type II pneumocyte | 0.2 | 0 | 0 | 0 | 1.8 |
| Edema/flooding, alveolar | 0 | 0 | 0 | 0 | 0.5 |
| Inflammation, bronchiolar | 1.2 | 0.4 | 0.4 | 0.2 | 2.4 |
| Hyperplasia, BALT | 1.2 | 0 | 0 | 0.4 | 0.1 |
| Infiltrates, granulocytes, peribronchiolar/bronchial | 0.2 | 0 | 0 | 0.8 | 0 |

**Figure 8.**
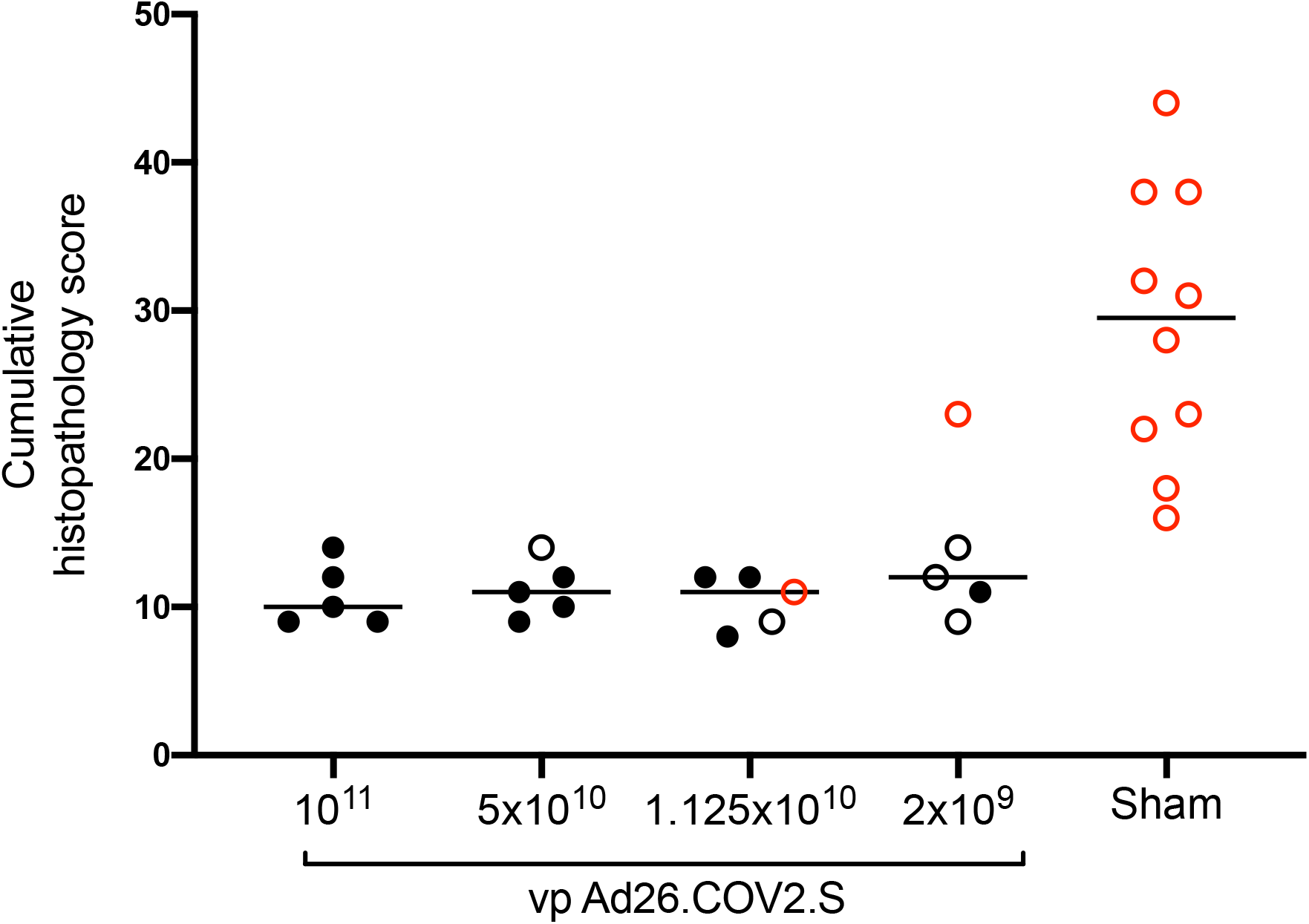
Histopathologic scoring of lung lesions in all lobes in vaccinated and sham animals following SARS-CoV-2 challenge. Scoring was performed independently by two blinded, veterinary pathologists. Lesions reported included 1) inflammation interstitial/septal thickening 2) infiltrate, macrophage 3) alveolar infiltrate, mononuclear 4) perivascular infiltrate, macrophage 5) bronchiolar type II pneumocyte hyperplasia 6) BALT hyperplasia 7) inflammation, bronchiolar/peribronchiolar infiltrate 8) neutrophils, bronchiolar/alveolar and 9) infiltrate, eosinophils. Lesions such as focal fibrosis and syncytia were reported but not included in scoring. Edema, alveolar flooding was excluded from scoring since animals received terminal bronchoalveolar lavages. Each feature assessed was assigned a score of 0= no significant findings; 1=minimal; 2= mild; 3=moderate; 4=marked/severe. Eight representative samples from cranial, middle, and caudal lung lobes from the left and right lungs were evaluated from each animal and were scored independently. Scores were added for all lesions across all lung lobes for each animal for a maximum possible score of 288 for each monkey. Lungs evaluated were inflated/suffused with 10% formalin. Horizontal lines reflect median values. Solid black circles indicate animals that showed no virus in BAL and NS following challenge, open black circles indicate animals that showed virus in NS but not BAL following challenge, and open red circles indicate animals that show virus in both BAL and NS following challenge.

## Discussion

In this study, we demonstrate that low doses of the Ad26.COV2.S vaccine protected rhesus macaques against SARS-CoV-2 challenge, although higher vaccine doses were required to protect in the upper respiratory tract as compared with the lower respiratory tract. Both activated memory B cells and binding and neutralizing antibody titers correlated with protective efficacy. Suboptimal vaccine dose levels led to viral breakthroughs in the upper respiratory tract, but no virologic, immunologic, histopathologic, or clinical evidence of VAERD was observed.

These data confirm and extend prior studies in which we showed that single-shot immunization with Ad26.COV2.S effectively protected against SARS-CoV-2 infection in rhesus macaques and against severe clinical disease in hamsters^5,6^. In the present study, we showed that Ad26.COV2.S doses as low as 2×10^9^ vp protected the lower respiratory tract, whereas doses of 1.125×10^10^ vp were required to protect the upper respiratory tract. This anatomic-specific difference in protective efficacy is consistent with multiple SARS-CoV-2 vaccine studies in nonhuman primates using DNA, RNA, and Ad vector-based vaccines, which have consistently shown superior protection in the lungs than in the nasal cavity^6-8,11^. These findings suggest that future work evaluating mucosal immune responses in these anatomic compartments is warranted.

Previous studies have suggested that vaccine-elicited binding and neutralizing antibodies correlated with protection against SARS-CoV-2 in rhesus macaques^6,11^ and that adoptive transfer of purified IgG from convalescent macaques protected against SARS-CoV-2 in this model^1^. The present data are consistent with these prior observations, and both RBD-specific activated memory B cells and binding and neutralizing antibody responses correlated with protective efficacy. Moreover, lower vaccine doses led to diminished antibody responses and reduced protective efficacy, further suggesting the importance of humoral immunity in protection against SARS-CoV-2. Suboptimal vaccine dose levels led to viral breakthroughs but did not result in evidence of enhanced viral replication or increased histopathologic pathology in the lungs of vaccinated animals compared with sham controls.

In summary, our data demonstrate that a single immunization of relatively low doses Ad26.COV2.S protects against SARS-CoV-2 challenge in rhesus macaques. Low dose vaccination led to viral breakthrough in the upper respiratory tract prior to the lower respiratory tract, raising the possibility that SARS-CoV-2 vaccines may protect against severe pneumonia more effectively than against asymptomatic upper respiratory tract infection in humans. Two phase 3 trials are currently in progress to determine the safety and efficacy of Ad26.COV2.S in humans.

## Acknowledgements

We thank M. Gebre, K. Verrington, E. Hoffman, L. Wrijil, T. Hayes, and K. Bauer for generous advice, assistance, and reagents. This project was funded in part by the Department of Health and Human Services Biomedical Advanced Research and Development Authority (BARDA) under contract HHS0100201700018C. We also acknowledge support from Janssen Vaccines & Prevention BV, Ragon Institute of MGH, MIT, and Harvard, Mark and Lisa Schwartz Foundation, Massachusetts Consortium on Pathogen Readiness (MassCPR), the National Institutes of Health (CA260476, OD024917, AI135098, AI129797, AI128751, AI126603, AI124377), and Fast Grants, Emergent Ventures, Mercatus Center at George Mason University.

## Author Contributions

D.H.B., R.Z., F.W., S.R.H., M.V.H., L.V.D.F., and H.S. designed the study and reviewed all data. X.H., A.C., J.Y., N.B.M., K.M., E.N.B., M.L., J.L., F.N., S.P., L.P., L.H.T. performed the immunologic and virologic assays. A.J.M., C.P.M., S.B., S.D., and R.C. performed the pathologic analyses. L.P., A.V.R., B.F., J.V., E.T., R.B., A.C., H.A., and M.G.L. led the clinical care of the animals. D.H.B. wrote the paper with all co-authors.

## Author Information

Correspondence and requests for materials should be addressed to D.H.B. D.H.B., R.Z., F.W. and H.S are co-inventors on provisional vaccine patents (63/121,482; 63/133,969; 63/135,182). R.Z., F.W., S.R.H., M.V.H., L.V.D.F., and H.S. are employees of Janssen Vaccines & Prevention BV and may hold stock in Johnson & Johnson.

## Data Availability Statement

All data are available in the manuscript and the supplementary material.

## Methods

### Animals and study design

30 outbred Indian-origin adult male and female rhesus macaques (*Macaca mulatta*) were randomly allocated to groups. All animals were housed at Bioqual, Inc. (Rockville, MD). Animals received a single immunization of 1×10^11^, 5×10^10^, 1.125×10^10^, or 2×10^9^ viral particles (vp) Ad26.COV2.S (Janssen; N=5/group) or sham (N=10) by the intramuscular route without adjuvant at week 0. At week 6, all animals were challenged with 1.0×10^5^ TCID50 (1.2×10^8^ RNA copies, 1.1×10^4^ PFU) SARS-CoV-2, which was derived from USA-WA1/2020 (NR-52281; BEI Resources) and deep sequenced^13^. Virus was administered as 1 ml by the intranasal (IN) route (0.5 ml in each nare) and 1 ml by the intratracheal (IT) route. All immunologic, virologic, and histopathologic studies were performed blinded. All animal studies were conducted in compliance with all relevant local, state, and federal regulations and were approved by the Bioqual Institutional Animal Care and Use Committee (IACUC).

### Subgenomic mRNA assay

SARS-CoV-2 E gene subgenomic mRNA (sgRNA or sgmRNA) was assessed by RT-PCR using primers and probes as previously described^11,13,17^. Briefly, to generate a standard curve, the SARS-CoV-2 E gene sgRNA was cloned into a pcDNA3.1 expression plasmid; this insert was transcribed using an AmpliCap-Max T7 High Yield Message Maker Kit (Cellscript) to obtain RNA for standards. Prior to RT-PCR, samples collected from challenged animals or standards were reverse-transcribed using Superscript III VILO (Invitrogen) according to the manufacturer’s instructions. A Taqman custom gene expression assay (ThermoFisher Scientific) was designed using the sequences targeting the E gene sgRNA^17^. Reactions were carried out on a QuantStudio 6 and 7 Flex Real-Time PCR System (Applied Biosystems) according to the manufacturer’s specifications. Standard curves were used to calculate sgRNA in copies per ml or per swab; the quantitative assay sensitivity was 50 copies per ml or per swab.

### Enzyme-linked immunosorbent assay (ELISA)

Binding antibodies were assessed by ELISA essentially as described^11,13^. Briefly, 96-well plates were coated with 1 μg/ml SARS-CoV-2 spike (S) or receptor binding domain (RBD) protein in 1X DPBS and incubated at 4°C overnight. After incubation, plates were washed once with wash buffer (0.05% Tween 20 in 1 X DPBS) and blocked with 350 μL Casein block/well for 2-3 h at room temperature. After incubation, block solution was discarded and plates were blotted dry. Serial dilutions of heat-inactivated serum diluted in casein block were added to wells and plates were incubated for 1 h at room temperature, prior to three further washes and a 1 h incubation with a 1:1000 dilution of anti-macaque IgG HRP (NIH NHP Reagent Program) at room temperature in the dark. Plates were then washed three times, and 100 μL of SeraCare KPL TMB SureBlue Start solution was added to each well; plate development was halted by the addition of 100 μL SeraCare KPL TMB Stop solution per well. The absorbance at 450nm was recorded using a VersaMax or Omega microplate reader. ELISA endpoint titers were defined as the highest reciprocal serum dilution that yielded an absorbance > 0.2. Log10 endpoint titers are reported.

### Pseudovirus neutralization assay

The SARS-CoV-2 pseudoviruses expressing a luciferase reporter gene were generated in an approach similar to as described previously^11-13^. Briefly, the packaging construct psPAX2 (AIDS Resource and Reagent Program), luciferase reporter plasmid pLenti-CMV Puro-Luc (Addgene), and spike protein expressing pcDNA3.1-SARS-CoV-2 SΔCT were co-transfected into HEK293T cells with calcium phosphate. The supernatants containing the pseudotype viruses were collected 48 h post-transfection; pseudotype viruses were purified by filtration with 0.45 μm filter. To determine the neutralization activity of the antisera from vaccinated animals, HEK293T-hACE2 cells were seeded in 96-well tissue culture plates at a density of 1.75 x 10^4^ cells/well overnight. Two-fold serial dilutions of heat inactivated serum samples were prepared and mixed with 50 μL of pseudovirus. The mixture was incubated at 37°C for 1 h before adding to HEK293T-hACE2 cells. After 48 h, cells were lysed in Steady-Glo Luciferase Assay (Promega) according to the manufacturer’s instructions. SARS-CoV-2 neutralization titers were defined as the sample dilution at which a 50% reduction in RLU was observed relative to the average of the virus control wells.

### IFN-γ enzyme-linked immunospot (ELISPOT) assay

ELISPOT plates were coated with mouse anti-human IFN-γ monoclonal antibody from BD Pharmingen at a concentration of 5 μg/well overnight at 4°C, and assays were performed as described^11,13^. Plates were washed with DPBS containing 0.25% Tween 20, and blocked with R10 media (RPMI with 11% FBS and 1.1% penicillin-streptomycin) for 1 h at 37°C. The Spike 1 and Spike 2 peptide pools contain 15 amino acid peptides overlapping by 11 amino acids that span the protein sequence and reflect the N- and C-terminal halves of the protein, respectively. Spike 1 and Spike 2 peptide pools were prepared at a concentration of 2 μg/well, and 200,000 cells/well were added. The peptides and cells were incubated for 18-24 h at 37°C. All steps following this incubation were performed at room temperature. The plates were washed with coulter buffer and incubated for 2 h with Rabbit polyclonal anti-human IFN-γ Biotin from U-Cytech (1 μg/mL). The plates are washed a second time and incubated for 2 h with Streptavidin-alkaline phosphatase antibody from Southern Biotechnology (1 μg/mL). The final wash was followed by the addition of Nitor-blue Tetrazolium Chloride/5-bromo-4-chloro 3 ‘indolyl phosphate p-toludine salt (NBT/BCIP chromagen) substrate solution for 7 min. The chromagen was discarded and the plates were washed with water and dried in a dim place for 24 h. Plates were scanned and counted on a Cellular Technologies Limited Immunospot Analyzer.

### IL-4 ELISPOT assay

Precoated monoclonal antibody IL-4 ELISPOT plates (Mabtech) were washed and blocked. The assay was then performed as described above except the development time with NBT/BCIP chromagen substrate solution was 12 min.

### B cell immunophenotyping

Fresh PBMCs were stained with Aqua live/dead dye for 20 min, washed with 2% FBS/DPBS buffer, and cells were suspended in 2% FBS/DPBS buffer with Fc Block (BD) for 10 min, followed by staining with monoclonal antibodies against CD45 (clone D058-1283, BUV805), CD3 (clone SP34.2, APC-Cy7), CD7 (clone M-T701, Alexa700), CD123 (clone 6H6, Alexa700), CD11c (clone 3.9, Alexa700), CD20 (clone 2H7, PE-Cy5), IgA (goat polyclonal antibodies, APC), IgG (clone G18-145, BUV737), IgM (clone G20-127, BUV396), IgD (goat polyclonal antibodies, PE), CD80 (clone L307.4, BV786), CD95 (clone DX2, BV711), CD27 (clone M-T271, BUV563), CD21 (clone B-ly4, BV605), CD14 (clone M5E2, BV570), CD138 (clone DL-101, PE-CF594), and staining with SARS-CoV-2 antigens including biotinylated SARS-CoV-2 RBD proteins (Sino Biological) and full-length SARS-CoV-2 spike proteins (Sino Biological) labeled with FITC and DyLight 405, at 4 °C for 30 min. After staining, cells were washed twice with 2% FBS/DPBS buffer, followed by incubation with BV650 streptavidin (BD Pharmingen) for 10min, then washed twice with 2% FBS/DPBS buffer. For intracellular staining, cells were permeabilized using Caltag Fix & Perm (Invitrogen), then stained with monoclonal antibodies against Ki67 (clone B56, PerCP-cy5.5) and IRF4 (clone 3E4, PE-Cy7). After staining, cells were washed and fixed by 2% paraformaldehyde. All data were acquired on a BD FACSymphony flow cytometer. Subsequent analyses were performed using FlowJo software (Treestar, v.9.9.6). For analyses, in singlet gate, dead cells were excluded by Aqua dye and CD45 was used as a positive inclusion gate for all leukocytes. Within class-switched B cell population gated as CD20+IgG+IgM-IgD-CD3-CD14-CD11c-CD123-CD7-, SARS-CoV-2 RBD-specific B cells were identified as double positive for SARS-CoV-2 RBD and spike proteins, and SARS-CoV-2 spike-specific B cells were identified as double positive for SARS-CoV-2 spike proteins labeled with different fluorescent probes. The SARS-CoV-2-specific B cells were further distinguished according to CD21/CD27 phenotype distribution: activated memory B cells (CD21-CD27+) and resting memory B cells (CD21+CD27+).

### Histopathology and immunohistochemistry

At time of fixation, lungs were suffused with 10% formalin to expand the alveoli. All tissues were fixed in 10% formalin and blocks sectioned at 5 μm. Slides were baked for 30-60 min at 65 degrees then deparaffinized in xylene and rehydrated through a series of graded ethanol to distilled water. Heat induced epitope retrieval (HIER) was performed using a pressure cooker on steam setting for 25 minutes in citrate buffer (Thermo; AP-9003-500) followed by treatment with 3% hydrogen peroxide. Slides were then rinsed in distilled water and protein blocked (BioCare, BE965H) for 15 min followed by rinses in 1x phosphate buffered saline. Primary rabbit anti-SARS-nucleoprotein antibody (Novus; NB100-56576) diluted 1:250, rabbit anti-Iba-1 antibody (Wako; 019-19741) diluted 1:250; rabbit anti-CD3 antibody (Sigma, SAB5500057) diluted 1:300, rabbit anti-CD20 (Invitrogen PA5-16701) diluted 1:750 followed by rabbit Mach-2 HRP-Polymer (BioCare; RHRP520L) for 30 minutes then counterstained with hematoxylin followed by bluing using 0.25% ammonia water. Labeling was performed on a Biocare IntelliPATH autostainer. All antibodies were incubated for 60 min at room temperature. Tissue pathology was assessed independently by two board-certified veterinary pathologists (AJM, RC).

### RNAscope

RNAscope in situ hybridization was performed as previously described^13^ using SARS-CoV2 anti-sense specific probe v-nCoV2019-S (ACD Cat. No. 848561) targeting the positive-sense viral RNA and DapB (ACD Cat.No 310043) as a negative control. In brief, after slides were deparaffinized in xylene and rehydrated through a series of graded ethanol to distilled water, retrieval was performed for 30 min in ACD P2 retrieval buffer (ACD Cat. No. 322000) at 95-98°C, followed by treatment with protease III (ACD Cat. No. 322337) diluted 1:10 in PBS for 20 min at 40°C. Slides were then incubated with 3% H2O2 in PBS for 10 minutes at room temperature. Slides were developed using the RNAscope® 2.5 HD Detection Reagents-RED (ACD Cat. No.322360).

### Statistical analyses

Analysis of virologic and immunologic data was performed using GraphPad Prism 8.4.2 (GraphPad Software). Comparison of data between groups was performed using two-sided Mann-Whitney tests. Correlations were assessed by two-sided Spearman rank-correlation tests. P-values of less than 0.05 were considered significant.

## References

1 McMahan, K. et al. Correlates of protection against SARS-CoV-2 in rhesus macaques. Nature, doi:10.1038/s41586-020-03041-6 (2020).

2 Abbink, P. et al. Comparative seroprevalence and immunogenicity of six rare serotype recombinant adenovirus vaccine vectors from subgroups B and D. J Virol 81, 4654–4663, doi:JVI.02696-06[pii]10.1128/JVI.02696-06 (2007).

3 Wrapp, D. et al. Cryo-EM structure of the 2019-nCoV spike in the prefusion conformation. Science 367, 1260–1263, doi:10.1126/science.abb2507 (2020).

4 Bos, R. et al. Ad26 vector-based COVID-19 vaccine encoding a prefusion-stabilized SARS-CoV-2 Spike immunogen induces potent humoral and cellular immune responses. NPJ Vaccines 5, 91, doi:10.1038/s41541-020-00243-x (2020).

5 Tostanoski, L. H. et al. Ad26 vaccine protects against SARS-CoV-2 severe clinical disease in hamsters. Nat Med 26, 1694–1700, doi:10.1038/s41591-020-1070-6 (2020).

6 Mercado, N. B. et al. Single-shot Ad26 vaccine protects against SARS-CoV-2 in rhesus macaques. Nature 586, 583–588, doi:10.1038/s41586-020-2607-z (2020).

7 Corbett, K. S. et al. Evaluation of the mRNA-1273 Vaccine against SARS-CoV-2 in Nonhuman Primates. N Engl J Med, doi: 10.1056/NEJMoa2024671 (2020).

8 van Doremalen, N. et al. ChAdOx1 nCoV-19 vaccine prevents SARS-CoV-2 pneumonia in rhesus macaques. Nature, doi:10.1038/s41586-020-2608-y (2020).

9 Gao, Q. et al. Development of an inactivated vaccine candidate for SARS-CoV-2. Science 369, 77–81, doi:10.1126/science.abc1932 (2020).

10 Wang, H. et al. Development of an Inactivated Vaccine Candidate, BBIBP-CorV, with Potent Protection against SARS-CoV-2. Cell 182, 713–721 e719, doi: 10.1016/j.cell.2020.06.008 (2020).

11 Yu, J. et al. DNA vaccine protection against SARS-CoV-2 in rhesus macaques. Science 369, 806–811, doi: 10.1126/science.abc6284 (2020).

12 Yang, Z. Y. et al. A DNA vaccine induces SARS coronavirus neutralization and protective immunity in mice. Nature 428, 561–564, doi:10.1038/nature02463 (2004).

13 Chandrashekar, A. et al. SARS-CoV-2 infection protects against rechallenge in rhesus macaques. Science 369, 812–817, doi:10.1126/science.abc4776 (2020).

14 Koutsakos, M. et al. Circulating TFH cells, serological memory, and tissue compartmentalization shape human influenza-specific B cell immunity. Sci Transl Med 10, doi:10.1126/scitranslmed.aan8405 (2018).

15 Neumann, B., Klippert, A., Raue, K., Sopper, S. & Stahl-Hennig, C. Characterization of B and plasma cells in blood, bone marrow, and secondary lymphoid organs of rhesus macaques by multicolor flow cytometry. J Leukoc Biol 97, 19–30, doi: 10.1189/jlb.1HI0514-243R (2015).

16 Titanji, K. et al. Acute depletion of activated memory B cells involves the PD-1 pathway in rapidly progressing SIV-infected macaques. J Clin Invest 120, 3878–3890, doi:10.1172/JCI43271 (2010).

17 Wolfel, R. et al. Virological assessment of hospitalized patients with COVID-2019. Nature, doi:10.1038/s41586-020-2196-x (2020).

