## Supplemental materials for "Low-Dose Ad26.COV2.S Protection Against SARS-CoV-2 Challenge in Rhesus Macaques"

### Supplementary Figure Legends

**Supplementary Figure S1. Study schema.**

**Supplementary Figure S2. B cell responses in vaccinated rhesus macaques.** Frequencies of RBD- and S-specific B cells in total IgG+ B cell populations following Ad26.COV2.S immunization. (**A**) Representative flow cytometry of PBMCs from one monkey in the $1x10^{11}$ vp dose group at days 0, 7, 14, and 28 after vaccination gated on class-switched IgG+ B cells. (**B**) Expression level of CD27 and CD95 on RBD-specific B cells. (**C**) Flow cytometry showing activated memory (AM) and resting memory (RM) B cells, gated on RBD-specific IgG+ B cells.

**Supplementary Figure S3. Correlations of B cell responses with adaptive immune responses.** Correlations of RBD-specific activated memory B cell frequencies with log NAb, log ELISA, and ELISPOT responses in vaccinated rhesus macaques. Red lines reflect the best linear fit relationship between these variables. P and R values reflect two-sided Spearman rank-correlation tests. n=25 biologically independent animals.

**Supplementary Figure S4. SARS-CoV-2 associated pathology in sham rhesus macaques following SARS-CoV-2 challenge.** Focal to locally extensive SARS CoV-2 associated pathological lesions were observed in sham vaccinated monkeys 10 days following challenge. (**A**) Bronchoepithelial syncytia (arrow, inset) within alveolus; (**B**) Multifocal Type II pneumocyte hyperplasia; (**C**) Inset from (**B**) showing type II pneumocyte hyperplasia (arrow) and endothelial reactivity (arrowhead); (**D**) Inset from (**C**) showing hyperplastic pneumocytes (arrow) and occasional polymorphonuclear cells (PMNs); (**E**) thrombus (arrow); (**F**) focal edema and consolidation due to pneumocyte hyperplasia; (**G**) multifocal interstitial pneumonia; (**H**) Inset from (**G**) showing large reactive cells. Lesions shown are from 4 animals. Scale bars = 20 microns (**G**), 50 microns (**A, C, E, F**), 200 microns (**B**).

**Supplementary Figure S5. Pathology in sham vaccinated animals corresponds to viral replication and inflammation following SARS-CoV-2 challenge.** (**A**) H&E showing type II pneumocyte hyperplasia; (**B**) Immunohistochemistry for SARS nucleocapsid protein; (**C**) RNAscope in situ hybridization for viral RNA (vRNA) in hyperplastic pneumocytes. Immunohistochemistry for (**D**) Iba-1 (macrophages), (**E**) CD3 (T lymphocytes) and (**F**) CD20 (B lymphocytes) in regions of lung pathology. All images from one representative sham animal 10 days following SARS-CoV-2 challenge. Scale bars = 20 microns (**A, B, D-F**), 50 microns (**C**).

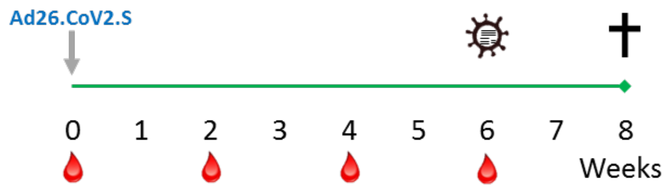


| Group | Ad26.COV2.S | N |
|---|---|---|
| 1 | $1x10^{11}$ vp | 5 |
| 2 | $5x10^{10}$ vp | 5 |
| 3 | $1.125x10^{10}$ vp | 5 |
| 4 | $2x10^{9}$ vp | 5 |
| 5 | N/A | 10 |

**Figure S1**

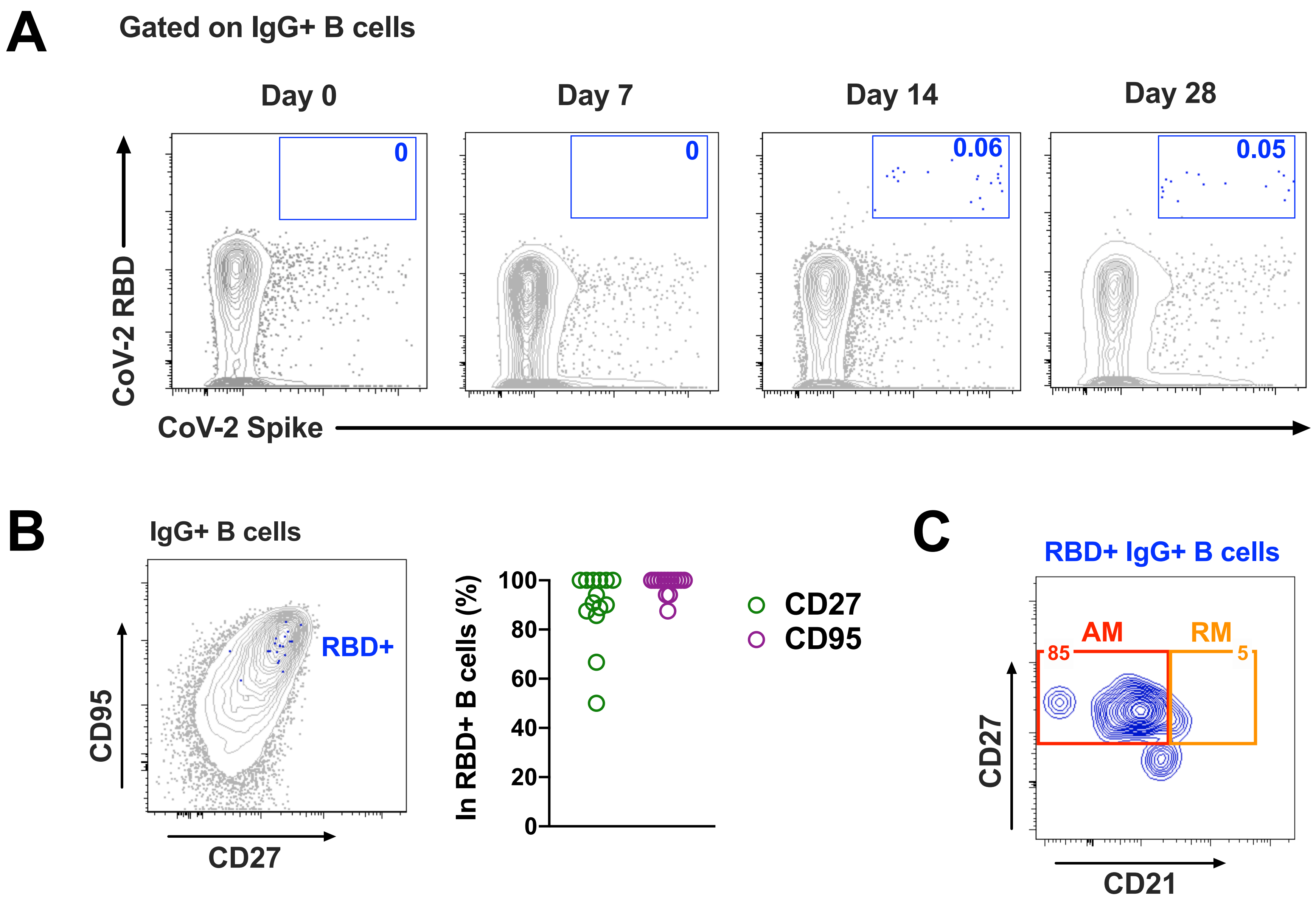
A
Gated on IgG+ B cells
Day 0
Day 7
Day 14
Day 28
0
0
0.06
0.05
CoV-2 RBD
CoV-2 Spike
B
IgG+ B cells
RBD+
CD95
CD27
In RBD+ B cells (%)
100
80
60
40
20
0
CD27
CD95
C
RBD+ IgG+ B cells
AM
RM
85
5
CD27
CD21

Figure S2

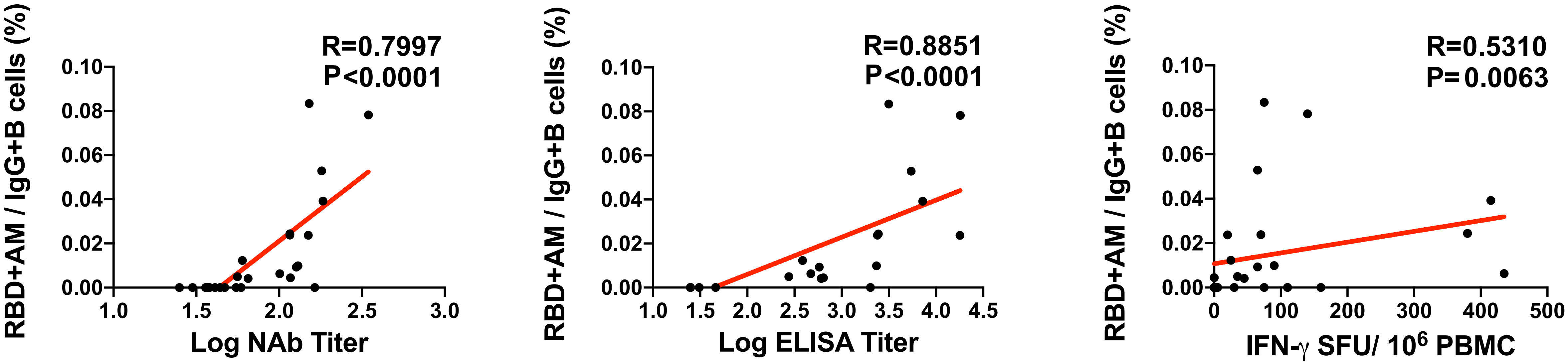
RBD+AM / IgG+B cells (%)
R=0.7997
P<0.0001
0.10
0.08
0.06
0.04
0.02
0.00
1.0
1.5
2.0
2.5
3.0
Log NAb Titer
RBD+AM / IgG+B cells (%)
R=0.8851
P<0.0001
1.0
1.5
2.0
2.5
3.0
3.5
4.0
4.5
Log ELISA Titer
RBD+AM / IgG+B cells (%)
R=0.5310
P= 0.0063
0
100
200
300
400
500
IFN-γ SFU/ 10⁶ PBMC

Figure S3

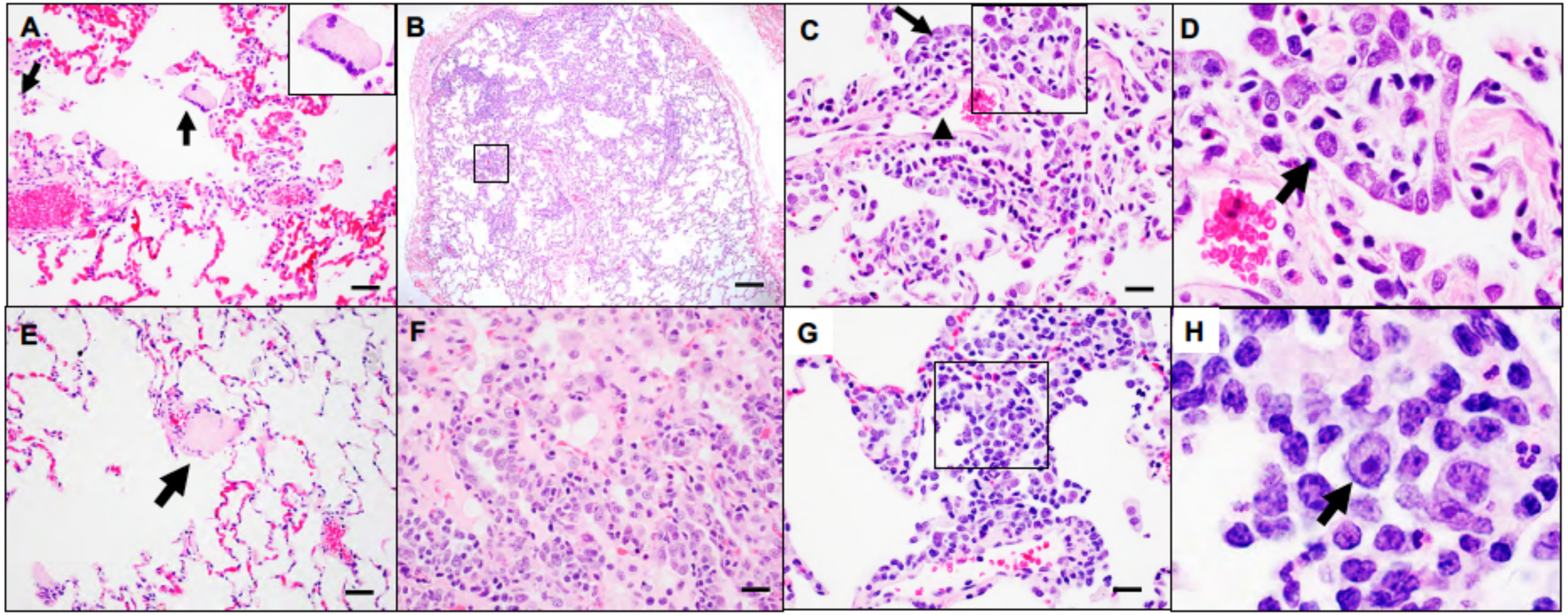


**Figure S4**

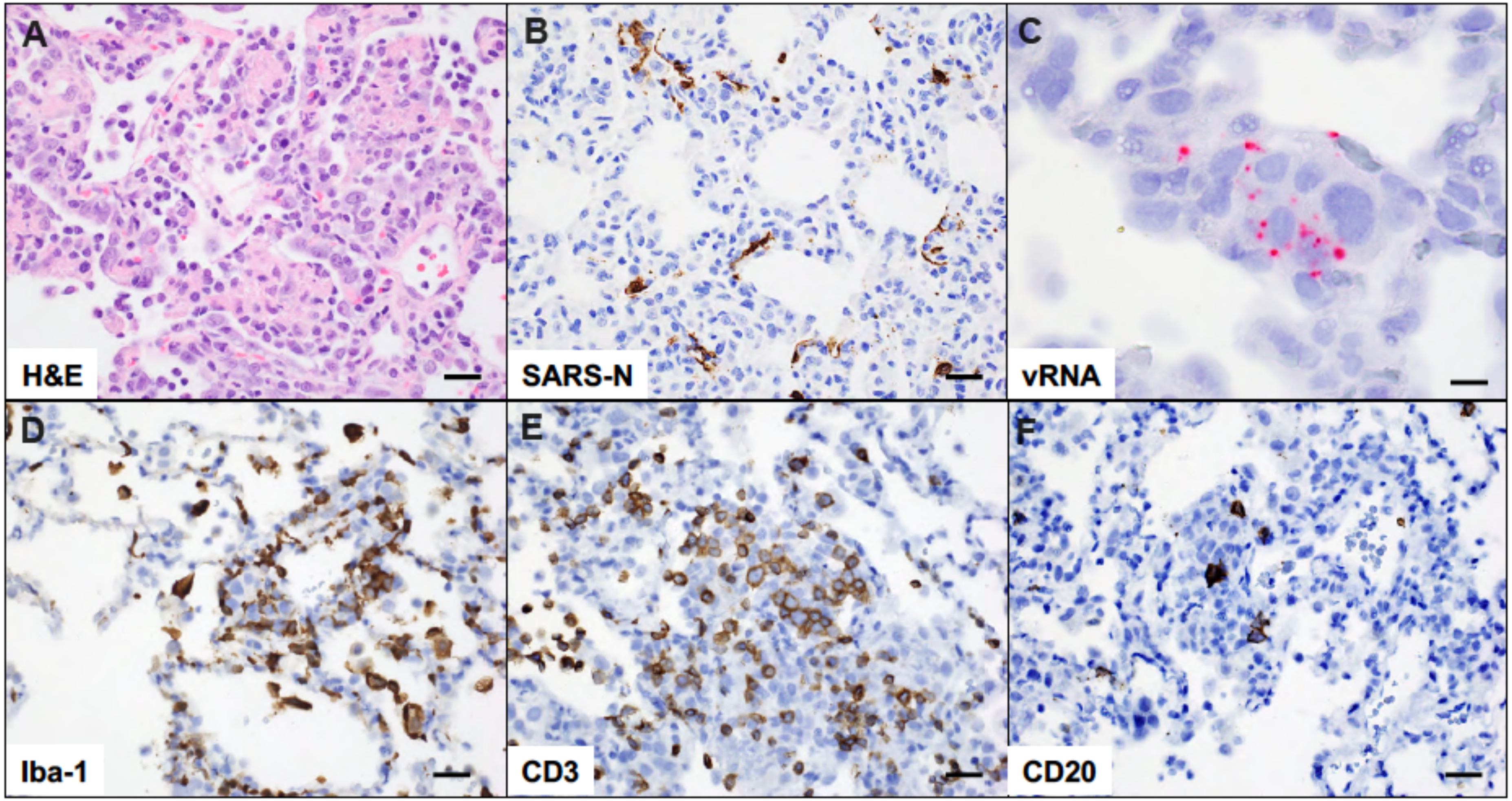


Figure S5
